# A comparison of resting state EEG and structural MRI for classifying Alzheimer’s disease and mild cognitive impairment

**DOI:** 10.1101/711465

**Authors:** FR Farina, DD Emek-Savaş, L Rueda-Delgado, R Boyle, H Kiiski, G Yener, R Whelan

**Author notes:** Correspondence to: Robert Whelan;, Francesca Farina.

## Abstract

Alzheimer’s disease (AD) is a neurodegenerative disorder characterised by severe cognitive decline and loss of autonomy. AD is the leading cause of dementia. AD is preceded by mild cognitive impairment (MCI). By 2050, 68% of new dementia cases will occur in low- and middle-income countries. In the absence of objective biomarkers, psychological assessments are typically used to diagnose MCI and AD. However, these require specialist training and rely on subjective judgements. The need for low-cost, accessible and objective tools to aid AD and MCI diagnosis is therefore crucial. Electroencephalography (EEG) has potential as one such tool: it is relatively inexpensive (cf. magnetic resonance imaging; MRI) and is portable. In this study, we collected resting state EEG, structural MRI and rich neuropsychological data from older adults (55+ years) with AD, with MCI and from healthy controls (n~60 per group). Our goal was to evaluate the utility of EEG, relative to MRI, for the classification of MCI and AD. We also assessed the performance of combined EEG and behavioural (Mini-Mental State Examination; MMSE) and structural MRI classification models. Resting state EEG classified AD and HC participants with moderate accuracy (AROC=0.76), with lower accuracy when distinguishing MCI from HC participants (AROC=0.67). The addition of EEG data to MMSE scores had no additional value compared to MMSE alone. Structural MRI out-performed EEG (AD vs HC, AD vs MCI: AROCs=1.00; HC vs MCI: AROC=0.73). Resting state EEG does not appear to be a suitable tool for classifying AD. However, EEG classification accuracy was comparable to structural MRI when distinguishing MCI from healthy aging, although neither were sufficiently accurate to have clinical utility. This is the first direct comparison of EEG and MRI as classification tools in AD and MCI participants.

## 1. Introduction

Alzheimer’s disease (AD) is a neurodegenerative disorder characterised by severe cognitive decline and loss of autonomy (i.e., dementia; Apostolova, 2016). AD is the leading cause of dementia, accounting for 70% of cases worldwide (Cassani, Estarellas, San-Martin, Fraga, & Falk, 2018). This number is expected to triple by 2050, with 68% of new dementia cases occurring in low- and middle-income countries (Prince et al., 2015). There is no known cure and only a handful of medications have been shown to delay symptom progression (Weller & Budson, 2018). Consequently, dementia risk reduction and prevention have been recognised as global priority areas for research (Shah et al., 2016).

Mild Cognitive Impairment (MCI) is a high-risk dementia condition marked by cognitive deficits that do not significantly impact daily living (Petersen & Negash, 2008). Approximately 3-15% of people with MCI are diagnosed with dementia every year (compared to 1-2% of the general population; Michaud, Su, Siahpush, Murman, et al., 2017). MCI therefore represents a promising avenue for early detection of AD. However, not all MCI diagnoses will be clinically relevant; 30% of people with MCI will not progress to AD within 6 years (Jicha et al., 2006). Early detection is also complicated by the fact that there is currently no inexpensive and practical diagnostic test for AD. Cerebrospinal fluid (CSF) and blood markers (e.g., Tau and Amyloid beta) have been identified, however, administering these tests is costly and not routine (Olsson et al., 2016). Instead, diagnosis is typically made using a combination of behavioural, neurological and psychological tests. These tests also have limitations; namely, they rely on subjective judgements, education level, and are time consuming and less sensitive at early stages of the disease (Cassani et al., 2018; Parra, 2014).

In recent years, considerable efforts have been made to identify neuroimaging biomarkers, with the aim of providing objective measures of disease pathology before symptom onset (Varghese, Sheelakumari, James, & Mathuranath, 2013). Most of this work has been carried out using structural Magnetic Resonance Imaging (MRI) (Park & Moon, 2016). Estimates of the sensitivity and specificity of MRI markers in detecting AD range from 82-89% and 86-97%, respectively (see Beheshti, Demirel, Matsuda, & Initiative, 2017, for a summary of recent studies). Detection rates are lower for MCI, with sensitivity estimated between 57-87% and specificity estimated between 52-82% (Beheshti et al., 2017). The most well-established structural MRI marker is medial temporal lobe (MTL) atrophy, including the hippocampus, entorhinal cortex and amygdala (Dubois et al., 2007). MTL atrophy is also the only neuroimaging marker included in the revised diagnostic criteria for AD (Jack Jr et al., 2018).

The practical utility of MRI as a diagnostic tool is restricted by high acquisition and infrastructure costs, and poor access to MRI scanners in many low- and middle-income countries (Musaeus et al., 2018). For example, in West Africa, a total of 84 MRI machines serve a combined population of over 350 million people (Ogbole, Adeyomoye, Badu-Peprah, Mensah, & Nzeh, 2018). Given the projected increase in global incidence of dementia, particularly in low- and middle-income countries, there is an urgent need for cheaper, more widely accessible tools to facilitate early diagnosis and disease management. Electroencephalography (EEG) represents one such potential alternative due to its low cost, portability and user-friendliness (i.e. not requiring participants to be immobile for long periods) (Cassani et al., 2018; Lizio et al., 2016).

EEG measures the summation of electrical dipoles created by, at least, thousands of cortical pyramidal neurons, and can therefore measure neurophysiological activation with millisecond precision. As such, EEG has the potential to detect neurophysiological signs of disease (e.g., abnormal synchronisation between brain regions) that may be present before any tissue loss occurs. Consequently, EEG markers may be more sensitive than structural MRI markers (which measures volume of neurons) at early stages in the disease process (Babiloni et al., 2016). Abnormal EEG patterns in dementia are well-documented (Tsolaki, Kazis, Kompatsiaris, Kosmidou, & Tsolaki, 2014). Typically, people with AD exhibit an increase in slow wave activity; that is, greater activity in lower frequency bands such as delta (1-4 Hz) and theta (4-8 Hz) and reduced activity in higher frequency bands such as alpha (8-13 Hz) and beta (13-30 Hz) (Musaeus et al., 2018). People with MCI also exhibit an increase in theta and a decrease in alpha power (Babiloni et al., 2006). Furthermore, increased theta and decreased (low) alpha (8-10 Hz) have been associated with conversion from MCI to AD (Moretti et al., 2009; Moretti et al., 2012).

Although EEG has potential to be a lower-cost alternative to MRI for AD diagnosis, its utility in dementia has yet to be firmly established. Results to date have been mixed; sensitivity and specificity estimates range from 0-90% and 65-88% in detecting AD, and from 55-77% and 66-100% in detecting MCI (e.g. Babiloni et al., 2016; Hatz et al., 2015; Lehmann et al., 2007; Musaeus et al., 2018). In addition to methodological differences (e.g., task-based or resting state, type of analysis, etc.), the generalisability of these findings is limited in many cases by small sample sizes (e.g., fewer than 15 patients) and potential confounding factors (Cassani et al., 2018). More specifically, Cassani and colleagues (2018) highlight that only 8 out of 112 studies investigating the use of EEG for AD diagnosis matched their patient and control groups for age, sex and education. This is particularly problematic as each of these variables has its own associated dementia risk (Prince et al., 2015).

In this study, we sought to identify the EEG features that best predict disease status by evaluating the efficacy of resting state (i.e., task-free) EEG in detecting AD, MCI and healthy aging. Features of interest included absolute power (i.e., summed power in different frequency bands), relative power (i.e., power in a given frequency band divided by the total power) and power ratios (i.e., power in one frequency band divided by another) (Cassani et al., 2018; Moretti et al., 2009). Power values were calculated separately for EEG recorded with participants’ eyes open and eyes closed to explore potential differences between these two arousal states (Barry & De Blasio, 2017). A secondary aim was to compare the accuracy of resting state EEG with that of structural MRI in classifying AD and MCI. EEG and MRI provide complementary information, which can inform our understanding of the neural mechanisms underlying cognitive decline (Moretti et al., 2012). However, to our knowledge, no study has evaluated both methods in AD and MCI participants. Finally, we assessed the utility of a combined behavioural-EEG model, using resting state EEG and Mini-Mental State Examination (MMSE), performance, as an additional low-cost option for classification. We predicted that structural MRI would out-perform resting state EEG in distinguishing AD participants from healthy controls, given the widespread grey matter loss associated with late-stage dementia, but that EEG (alone or in combination with behavioural performance) would be better at distinguishing MCI participants from healthy participants. We further hypothesised that delta and theta power would be the best predictors of AD status, and that theta power would be the best predictor of MCI status.

## 2. Methods

### 2.1. Study population

Community-dwelling older adults with AD (n = 118) and those with amnestic MCI (n = 134) were recruited from the outpatient memory clinic of the Department of Neurology at Dokuz Eylul University. Healthy older adults (n = 198; aged 55 years and above) were recruited from various community sources including announcements in public conferences and university billboards. Diagnosis of AD was made according to the National Institute on Aging and Alzheimer’s Association (NIA-AA) criteria (McKhann et al., 2011). Inclusion criteria for participants with AD were: a) insidious onset; b) impairment of daily functioning (Clinical Dementia Rating (CDR) score of ≥1); c) impairment of two or more cognitive domains; and d) exclusion of delirium, other causes of dementia and other major psychiatric disorders. Diagnosis of MCI was also made according to the NIA-AA criteria (Albert et al., 2011). Inclusion criteria for participants with MCI were: a) objective memory impairment (1-1.5 standard deviations below the mean) on verbal and visual memory tests (see 2.2. Diagnostic Criteria); b) normal activities of daily living documented by history and their relatives; and c) a CDR score of 0.5. All participants with MCI and AD underwent MRI and laboratory tests to rule out other causes of cognitive impairment.

Exclusion criteria for participants with MCI and AD were: a) history of depression or psychosis; b) history or signs of major stroke; c) presence of any other neurological and/or psychiatric diseases including drug addiction, alcohol abuse or epilepsy; d) uncontrolled systemic diseases and e) traumatic brain injury. Inclusion criteria for healthy control (HC) participants were: a) no neurological abnormality or global cognitive impairment (Mini-Mental State Examination (MMSE) score ≥27); b) no history of psychiatric or systemic disorders; and c) no history of severe head injury and alcohol or drug abuse. All participants had normal or corrected-normal vision. The study was approved by the Department of Neurology Institutional Review Board at Dokuz Eylul University. All participants provided informed consent according to the guidelines outlined in the Declaration of Helsinki.

### 2.2. Diagnostic criteria

Neurological, neuroimaging (MRI) and laboratory examinations were carried out. All participants were assessed using a comprehensive battery of neuropsychological tests designed to assess verbal and visual episodic memory, attention, executive functions, visuospatial skills and language using the following tests: MMSE (Folstein, Folstein, & McHugh, 1975), Öktem Verbal Memory Processes Test (OVMPT) (Öktem, 1992), Wechsler Memory Scale-Revised (WMS-R) Digit Span Test (Wechsler, 1987), Verbal Fluency Test (semantic) (Tumac, 1997), Boston Naming Test (BNT) (Kaplan, Goodglass, & Weintraub, 2001; Mack, Freed, Williams, & Henderson, 1992) and CDR Scale (Hughes, Berg, Danziger, Coben, & Martin, 1982). The Yesavage Geriatric Depression Scale was also administered to participants to test the exclusion criterion for depression (Yesavage et al., 1982).

### 2.3. EEG data acquisition

EEGs were recorded from 30 Ag/AgCl electrodes positioned on an elastic cap (Easy-Cap; Brain Products GmbH; Gilching, Germany) according to the international 10-20 system were referenced to linked earlobe electrodes (A1 + A2). The recording room was electrically shielded, sound attenuated and dimly illuminated. The electrooculogram (EOG) was registered from both the medial upper and the lateral orbital rim of the right eye. All electrode impedances were kept less than 10 kΩ. EEG and EOG were amplified by means of a Brain Amp 32-channel DC system machine with 0.03–70 Hz bandpass filter and were digitized online with a sampling rate of 500 Hz (Brain Products GmbH; Gilching, Germany). EEGs were recorded eyes open (EO) for four minutes and eyes closed (EC) for four minutes.

### 2.4. EEG data pre-processing

EEG data were pre-processed using the EEGLAB toolbox (Delorme & Makeig, 2004) in conjunction with the FASTER plug-in (Fully Automated Statistical Thresholding for EEG artefact Rejection; Nolan, Whelan, & Reilly, 2010). Data were bandpass filtered between 0.1 and 70 Hz, notch filtered at 50 Hz and average referenced across all scalp electrodes. Data were then epoched into 2-second segments. FASTER removed epochs containing large artefacts (e.g. muscle twitches) and interpolated channels with poor signal quality. Artefactual (i.e., non-neural) independent components were also identified and removed from the data automatically using FASTER. All pre-processing parameters for FASTER for this study are included in the Supplemental Material (Supplementary material_FASTER_processing.eegjob). Data were then visually inspected for quality and to remove any remaining noisy data.

### 2.5. EEG frequency band and power ratio calculations

Spectral analysis of absolute and relative power across the 30 scalp electrodes was conducted using the multitaper spectral estimation with Hanning taper and 0.5 frequency resolution. The following seven frequency bands were included: delta (1-4 Hz), theta (4-8 Hz), alpha1 (8-10 Hz), alpha2 (10-13 Hz), beta1 (13-18 Hz), beta2 (18-30 Hz) and gamma (30-45 Hz), based on previous literature (Cassini et al., 2018). For absolute power, raw values were log scaled to produce values in dB. For relative power, the values were expressed as a percentage of power in a frequency band divided by the total power across all seven frequency bands. Three additional power metrics were also calculated for the EO condition, based on previous literature (Moretti et al., 2009; Moretti et al., 2012; Musaeus et al., 2018). These were: theta/gamma power ratio, alpha2/alpha1 power ratio and global theta power. Theta/gamma ratios were calculated by dividing the absolute power in the theta band with the absolute power in the gamma band to create a ratio for each electrode. Alpha2/alpha1 ratios were calculated in the same way. Global theta was calculated by averaging absolute theta power for all channels. Absolute and relative power were calculated separately for EO and EC conditions.

### 2.6. MRI data acquisition

MRI data were acquired using a 1.5 Tesla Achieva MRI scanner (Philips Medical Systems, Best, The Netherlands). Participants underwent a 10-min T1 scan as part of a larger 20-min MRI battery. Two separate protocols were used for scans: the Alzheimer’s Disease Neuroimaging Initiative (ADNI) T1 protocol (129 scans) and a local protocol (49 scans). The ADNI protocol used the turbo field echo sequence with the following parameters: number of slices = 166, FOV = 240 mm^3^, slice thickness = 1 mm, slice gap = 0 mm, TR = 9 ms, TE = 4 ms. The local protocol used a gradient echo sequence with the following parameters: number of slices = 150, FOV = 230 mm^3^, slice thickness = 1 mm, slice gap = 0 mm, TR = 25 ms, TE = 6 ms.

### 2.7. MRI data pre-processing

Before processing, scans were automatically reoriented to a canonical SPM template and visually inspected for good orientation and gross artefacts. Poorly oriented scans were manually reoriented. Images were then pre-processed using SPM12 (University College London, London, UK), separately for each acquisition protocol. Bias correction was applied, and images were segmented into grey matter, white matter and cerebrospinal fluid. SPM DARTEL was used for non-linear registration of the segmented grey matter images to a custom template. Images were affine registered to MNI space and resampled (1 mm^3^) with modulation to preserve the signal from each voxel. Images were then smoothed with a 4 mm full-width at half maximum Gaussian kernel and visually inspected for accurate segmentation (full pre-processing information is available at github.com/rorytboyle/brainPAD).

### 2.8. Machine learning

A machine learning analysis with penalized logistic regression was used to estimate the accuracy of (1) resting state EEG and (2) structural MRI data in classifying individuals as AD, MCI or HC. Binary classification was performed with (1) AD vs HC participants, (2) MCI vs HC participants and (3) AD vs MCI participants. As a control analysis, we also calculated the diagnostic accuracy of MMSE data using the same method. The MMSE is the most widely used test to measure cognitive ability for AD diagnosis (Cassani et al., 2018). Hence, we included the MMSE analysis to provide ‘ceiling’ classification rates in our sample, against which to compare the results of the neuroimaging models (cf. Rossini et al., 2008). However, it should be noted that the MMSE is less sensitive at detecting MCI (Nasreddine et al., 2005), and so we did not expect accuracy to be as high for classifying individuals with MCI compared to AD.

#### 2.8.1. Data preparation

In preparation for analysis, AD, MCI and HC groups were matched for age, sex and years of education (see Table 1), all of which are confounding factors in AD (Cassani et al., 2018). Group matching was done separately for MMSE, EEG and MRI datasets to ensure that the groups were matched across all modalities (see Figure S1). Group sizes were also approximately matched to minimise any bias toward one particular group in the analysis (cf. Raamana, Weiner, Wang, Beg, & Alzheimer’s Disease Neuroimaging Initiative, 2015). For the MMSE data, the final matched sample consisted of 180 participants (57 AD, 61 MCI and 62 HC). For the EEG data, the sample consisted of 189 participants (60 AD, 64 MCI and 65 HC). For the MRI data, the sample consisted of 178 participants (60 AD, 60 MCI and 58 HC).

**Table 1.** Demographic information of AD, MCI and HC groups included in the MMSE, EEG and MRI classification analyses, and results of their statistical comparisons.

| Measure |  | AD | MCI | HC | Stat. test (df) | P |
| --- | --- | --- | --- | --- | --- | --- |
| MMSE | *n* | 57 | 61 | 62 | - | - |
| | Age | 71.30 (7.14) | 71.23 (4.39) | 69.95 (4.85) | $F(177) = 1.13$ | 0.33 |
| | Sex, F (%) | 34 (59.65) | 30 (48.39) | 37 (60.66) | $\chi^2(177) = 2.29$ | 0.32 |
| | Education | 9.37 (4.35) | 10.07 (4.75) | 9.41 (5.16) | $F(177) = 0.41$ | 0.67 |
| EEG | *n* | 60 | 64 | 65 | - | - |
| | Age | 71.20 (7.07) | 71.31 (4.34) | 70.06 (4.79) | $F(186) = 1.02$ | 0.36 |
| | Sex, F (%) | 36 (60) | 31 (48.44) | 41 (63.08) | $\chi^2(186) = 3.10$ | 0.21 |
| | Education | 9.35 (4.29) | 9.98 (4.88) | 9.15 (5.16) | $F(186) = 0.52$ | 0.59 |
| MRI | *n* | 60 | 60 | 58 | - | - |
| | Age | 70.85 (9.59) | 71.85 (5.43) | 69.71 (6.62) | $F(175) = 1.23$ | 0.30 |
| | Sex, F (%) | 38 (63.33) | 31 (51.67) | 38 (65.52) | $\chi^2(175) = 2.74$ | 0.26 |
| | Education | 9.07 (4.32) | 9.85 (4.49) | 10.72 (4.95) | $F(175) = 1.92$ | 0.15 |

Values are mean (standard deviation) or number (percentage). One-way between-groups analysis of variance (ANOVA) was used for all group comparisons, except for sex, which was assessed via Kruskal-Wallis test. Note: Demographic information for the MMSE+EEG models is the same as the MMSE models.

Thirty separate classification analyses were run. This included three MMSE models (AD vs HC, MCI vs HC, AD vs MCI), 21 EEG models (absolute power EO, absolute power EC, relative power EO, relative power EC, theta/gamma ratio, alpha2/alpha1 ratio and global theta power for each group comparison), three MRI models (AD vs HC, MCI vs HC and AD vs MCI), and three MMSE+EEG models (AD vs HC, MCI vs HC and AD vs MCI). For all models, the model input was a data matrix of participants (rows) x features (columns). MMSE models contained a single feature per participant (i.e., MMSE scores). Absolute power EO, absolute power EC, relative power EO and relative power EC models contained 210 features per participant (30 channels x 7 frequency bands). Each feature represented activation in a specific scalp location for one frequency band. Theta/gamma and alpha2/alpha1 ratio models contained 30 features each (i.e., 30 channels x 1 ratio value). Global theta models consisted of a single feature (i.e., summed theta power). MRI models consisted of 2,735 features per participant, each representing a specific anatomical location in 3D space. MMSE+EEG models contained two features (i.e. MMSE scores + summed theta power).

#### 2.8.2. Classification method

The same classification method was used for all models. This method has previously been reported in detail by Kiiski and colleagues (2018). A brief description is provided here (see also Figure S2). One important modification regarding cross-validation should be noted. Data in the current study were divided into five cross-validation folds as opposed to 10 folds (per Kiiski et al., 2018). As such, each model was trained on 80% of the sample (the training set) and applied to the remaining 20% (the test set). This modification was designed to by including more data in the test set, while also reserving enough training data to achieve good model fit (Varoquaux et al., 2018). Per Kiiski *et al*., cross-validation was also applied within each training set (known as ‘nested’ cross-validation) to tune model hyperparameters. This was done by further subdividing each main fold into nested training (64% of the total data) and nested test sets (16% of the total data). Parameters that yielded the lowest prediction error at the sub-fold level were used to fit a model using the training set of the main fold.

To further quantify model performance, the entire analysis was repeated 100 times with different allocations of participants to training and testing folds, and the results aggregated across the 100 iterations. The cross-validated result was then compared to a ‘null’ model to quantify the baseline classification level. The null model was generated by random-label permutation, whereby individuals were assigned randomly to an outcome measure from another participant, and the entire machine learning analysis was repeated. The accuracy of the null model was then compared to the accuracy of the actual model (i.e., with real data) using a t-test. The actual model was considered successful if the Area Under the Curve (AUC) value was statistically significantly higher than that of the null model (*p* < 0.0016, 0.05/30; corrected for multiple comparisons). Three additional indices of model performance were also used for model interpretation. These were (1) *accuracy* (i.e. the number of true positive (TP) and true negative (TN) cases divided by the sample size), (2) *sensitivity* (i.e. the number of TP values correctly identified) and (3) *specificity* (i.e. the number of TN values identified as negative).

To identify the most predictive features in successful models, we computed the ‘selection frequency’ of each individual feature (cf. Rueda-Delgado et al., 2019). Selection frequency was calculated by summing each feature’s non-zero count in each main fold (i.e., the number of times that feature was selected in the model) and then averaging this value across the 100 repetitions. Features selected in ≥90% of models were deemed to be robustly predictive.

## 3. Results

### 3.1. Demographic information

Table 1 summarises the demographic information of the matched groups included in the MMSE, EEG and MRI classification models, along with results of their statistical comparisons.

### 3.2. Cognitive performance

A small number of values were missing from the OVMPT (2 AD), Digit Span test (3 AD, 6 MCI), Verbal Fluency test (1 AD, 1 HC), BNT (1 AD, 2 MCI, 4 HC) and MMSE (3 AD, 2 MCI, 4 HC). Table 2 summarises the remaining data for the groups included in the MMSE, EEG and MRI models. Differences were assessed using one-way between-groups ANOVA followed by Bonferroni-corrected pairwise comparisons. Bayesian ANOVA were also carried out for non-significant effects. Group differences were found for all behavioural tests in the MMSE, EEG and MRI sub-samples. HC groups performed significantly better than MCI and AD groups on the MMSE (all *p*s < 0.001), OVMPT Total Learning (all *p*s < 0.002) and Delayed Recall tests (all *p*s < 0.001) and the Verbal Fluency test (all *p*s < 0.001). MCI groups performed better than AD groups on the MMSE, OVMPT Total Learning and Delayed Recall tests and the Verbal Fluency test (all *p*s < 0.001). HC and MCI groups also out-performed AD groups on the Digit Span Backward (all *p*s < 0.04) and the BNT (all *p*s < 0.002). No significant differences were found between the HC and MCI groups in these tests (all *p*s > 0.08). The majority of Bayes Factors (BF) computed for non-significant results provided anecdotal evidence in the direction of either hypothesis (BF10 values between 0.25-1.63). However, moderate to strong support for a difference between MCI and HC performance on the BNT in the MRI sample was found (BF_10_ = 0.08).

**Table 2.** Neuropsychological test performance for all groups and results of their statistical comparisons.

|  | **Measure** | **AD** | **MCI** | **HC** | **Stat. test (df)** | **P value** | **Sig. comparisons** |
| --- | --- | --- | --- | --- | --- | --- | --- |
| MMSE | Mini-Mental State Examination | 20.95 (4.17) | 26.46 (2.87) | 28.62 (1.58) | $F(164) = 94.88$ | <0.001 | All groups |
| | OVMPT Total Learning | 50.71 (18.21) | 75.43 (17.56) | 113.76 (11.60) | $F(164) = 223.71$ | <0.001 | All groups |
| | OVMPT Delayed Recall | 2.02 (2.71) | 6.83 (3.37) | 13.38 (1.36) | $F(164) = 273.41$ | <0.001 | All groups |
| | Verbal Fluency | 12.55 (4.96) | 17.09 (4.82) | 21.02 (4.77) | $F(164) = 43.13$ | <0.001 | All groups |
| | Digit Span Backward | 2.91 (1.21) | 3.43 (0.82) | 3.78 (1.19) | $F(164) = 9.05$ | <0.001 | AD vs HC, AD vs MCI |
| | Boston Naming Test | 13.33 (1.96) | 14.39 (1.27) | 14.79 (0.49) | $F(164) = 17.17$ | <0.001 | AD vs HC, AD vs MCI |
| EEG | Mini-Mental State Examination | 20.72 (4.31) | 26.24 (2.88) | 28.61 (1.55) | $F(177) = 100.35$ | <0.001 | All groups |
| | OVMPT Total Learning | 51.16 (17.84) | 74.94 (18.72) | 112.25 (12.34) | $F(184) = 216.3$ | <0.001 | All groups |
| | OVMPT Delayed Recall | 1.95 (2.66) | 6.75 (3.63) | 13.22 (1.44) | $F(184) = 266.25$ | <0.001 | All groups |
| | Verbal fluency | 12.66 (4.98) | 17.11 (4.59) | 20.69 (4.74) | $F(184) = 43.65$ | <0.001 | All groups |
| | Digit Span Backward | 2.92 (1.18) | 3.43 (0.80) | 3.72 (1.17) | $F(177) = 8.44$ | <0.001 | AD vs HC, AD vs MCI |
| | Boston Naming Test | 13.34 (1.90) | 14.39 (1.21) | 14.75 (0.54) | $F(179) = 18.31$ | <0.001 | AD vs HC, AD vs MCI |
| MRI | Mini-Mental State Examination | 19.60 (4.73) | 25.79 (2.80) | 28.72 (1.41) | $F(173) = 114.83$ | <0.001 | All groups |
| | OVMPT Total Learning | 45.03 (16.14) | 71.22 (15.72) | 113.79 (12.60) | $F(173) = 316.50$ | <0.001 | All groups |
| | OVMPT Delayed Recall | 1.47 (2.46) | 5.97 (3.13) | 13.02 (1.36) | $F(174) = 338.03$ | <0.001 | All groups |
| | Verbal fluency | 11.16 (5.08) | 16.43 (4.41) | 21.70 (4.59) | $F(174) = 72.05$ | <0.001 | All groups |
| | Digit Span Backward | 2.71 (1.38) | 3.53 (0.95) | 4.02 (1.01) | $F(166) = 19.57$ | <0.001 | AD vs HC, AD vs MCI |
| | Boston Naming Test | 13.11 (2.22) | 14.14 (1.53) | 14.80 (0.49) | $F(172) = 16.41$ | <0.001 | AD vs HC, AD vs MCI |

Values are mean (standard deviation) or number (percentage). Significant group comparisons indicate $p < 0.05$ .

### 3.3. Classification results

Figure 1 shows results from the MMSE, EEG, MRI and MMSE+EEG models, including AUC, accuracy, sensitivity and specificity. A full description of the features selected in each model is provided in the Supplementary material (Model_features.xls).

**Figure 1.**
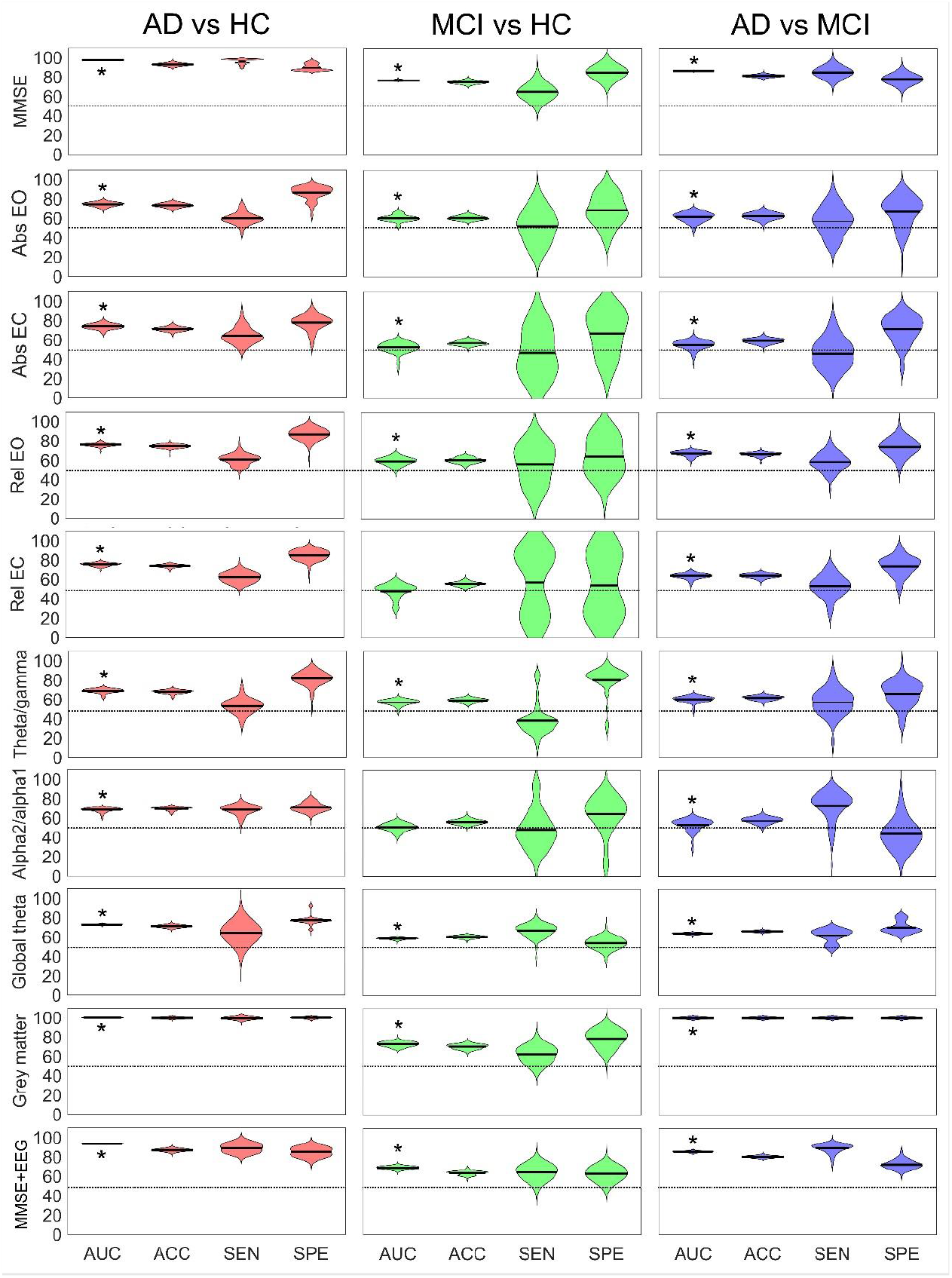
Classification results for the MMSE, EEG, MRI and EEG+MRI models comparing AD vs HC, MCI vs HC and AD vs MCI groups. Prediction indices of the MMSE, EEG and MRI models over 100 repetitions. Indices include accuracy (ACC, i.e., number of true positives and true negatives divided by the sample size), Area Under the Curve (AUC), Sensitivity (SEN; i.e. true positive rate) and Specificity (SPE, i.e., true negative rate). AUC values significantly higher than the null models are indicated by * (*p* < 0.001). Dashed line indicates 50% (i.e. chance level). Abbreviations: Abs, absolute power; EC, eyes closed; EO, eyes open; Rel, relative power.

#### 3.3.1. MMSE models

Figure 2 illustrates Receiver Operating Curves (ROCs) for the MMSE models. The AD vs HC model resulted in the best performance (AUC = 0.97). This was followed by the AD vs MCI model (AUC = 0.87) and then the MCI vs HC model (AUC = 0.77). Prediction indices were high overall, with the exception of sensitivity in the MCI vs HC model, which was only moderately good (65%). All models significantly out-performed 100% of null model iterations (all *t*s > 91, all *p*s < 0.001). MMSE scores were predictive of HC status in the AD and MCI models, and MCI status in the AD vs MCI model.

**Figure 2.**
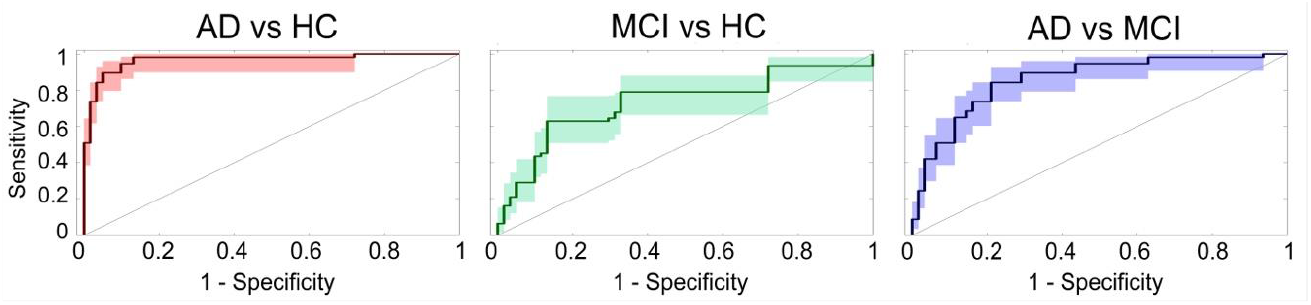
Receiver Operating Curves (ROCs) for MMSE actual models with 95% confidence intervals. Curves close to the top left corner (0.0, 1.0) indicate good performance.

#### 3.3.2. EEG models

Data from 42 participants (16 AD, 13 MCI and 13 HC) were excluded due to excessive noise. The remaining sample consisted of 408 participants (102 AD, 121 MCI and 185 HC). The total percentage of data removed during pre-processing and visual inspection was 6.38% (± 2.78) for the EC condition and 6.67% (± 5.91) for the EO condition. Mean duration of the final data was 225 s (± 9.89 s) for EC and 224 s (± 12.72 s) for EO.

Figure 3 illustrates ROCs for the EEG models. For AD vs HC classification, this was relative power EC (AUC = 0.76). Performance was also good for relative power EO, absolute power EC and EO, and global theta models (AUCs > 0.70). Theta/gamma and alpha2/alpha1 ratio models yielded moderate results (AUC = 0.69 and 0.68, respectively). AD vs HC models had high specificity (i.e. 70-87%), but only weak to moderate sensitivity (i.e. 53-68%). All models significantly out-performed >99% of null models (all *t*s > 38, all *p*s < 0.001). Theta power was the best predictor of AD status across models, including both absolute and relative power EO and EC. Theta in multiple frontal, central, temporal and parietal areas survived the selection threshold (see Figure 4A). The best predictors of HC status were left temporo-parietal alpha2/alpha1 ratio and beta1 power in the relative power EO model (see Figure 4A).

**Figure 3.**
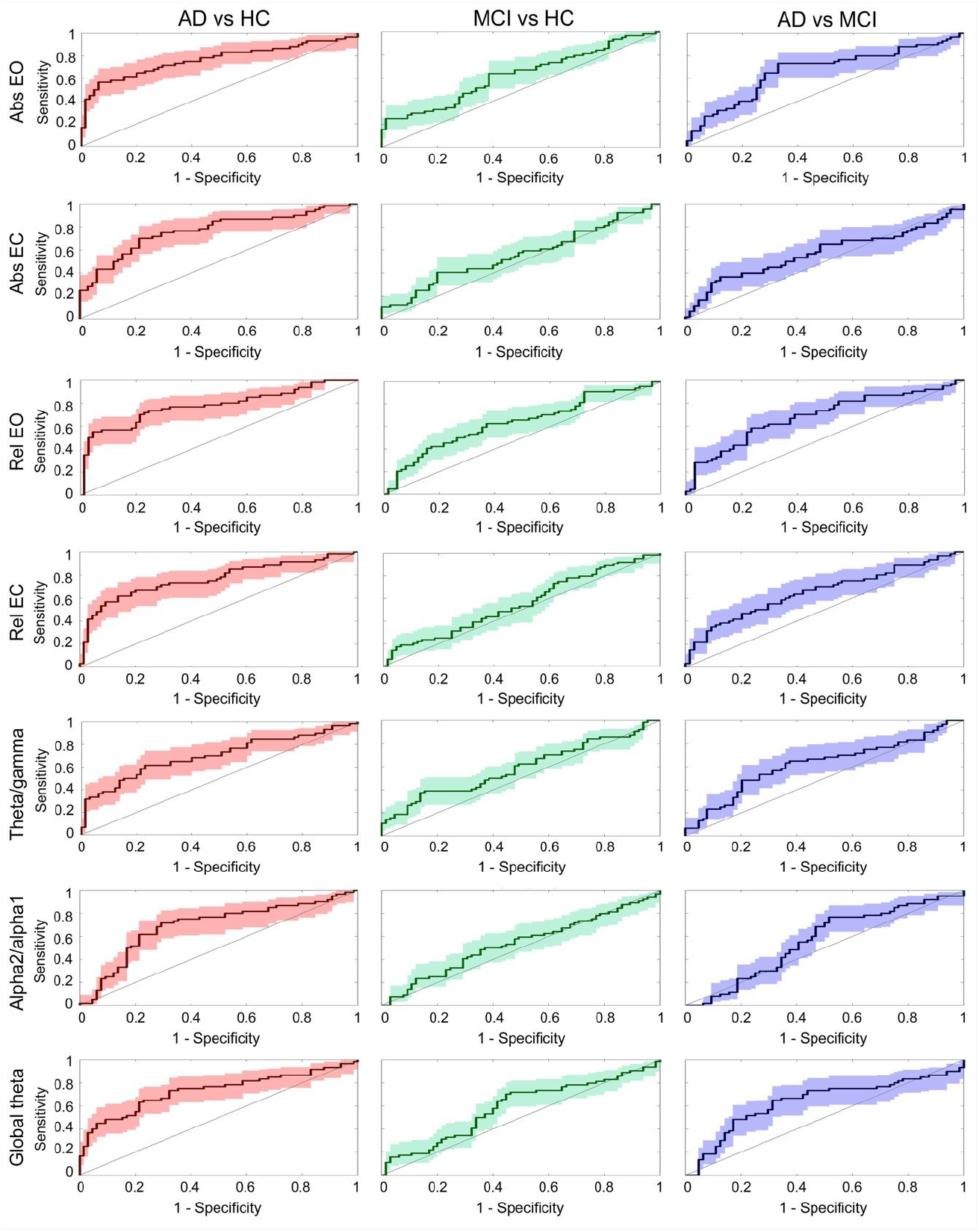
Receiver Operating Curves (ROCs) for EEG models with 95% confidence intervals. Curves close to the top left corner (0.0, 1.0) indicate good performance. Abbreviations: Abs, absolute power; EC, eyes closed; EO, eyes open; Rel, relative power.

**Figure 4.**
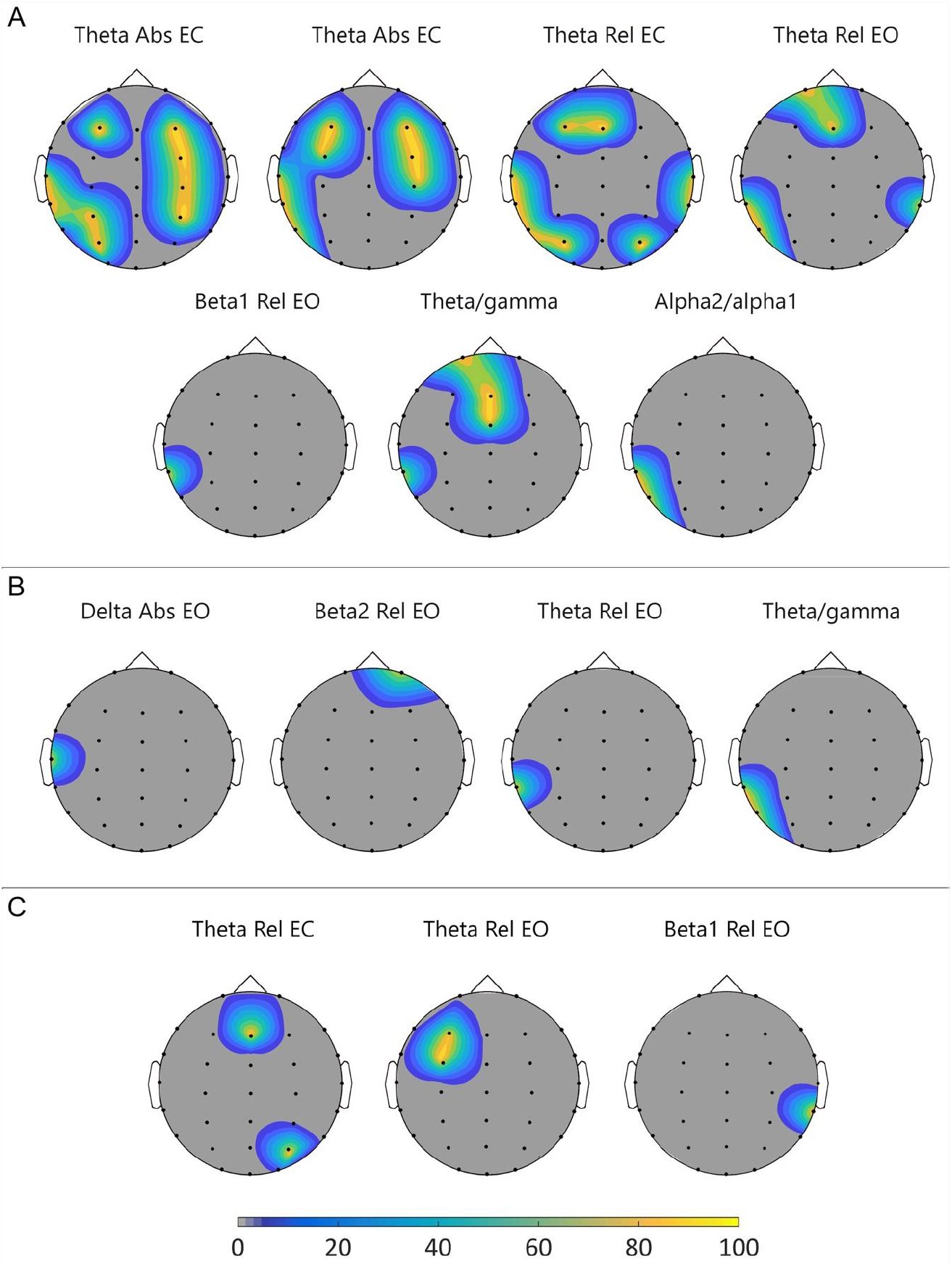
Topographic plots of the selection frequency at selected electrodes for (A) AD vs HC, (B) MCI vs HC and (C) AD vs MCI models. The colour represents the selection frequency of the features across iterations that survived the 90% threshold. Abbreviations: Abs: Absolute power; EC: eyes closed; EO: eyes open; Rel: Relative power.

The best model for MCI vs HC classification was absolute power EO (AUC = 0.61), which out-performed 94% of null model iterations (*t*(198) = 20.90, *p* < 0.001). All other models produced weaker effects (AUCs < 0.60). Absolute power EC, relative power EO, theta/gamma ratio and global theta power models significantly out-performed >66% of null model iterations (all *t*s > 4, all *p*s < 0.001). Alpha2/alpha1 ratio and relative power EC models did not perform better than the null model (*t*s < 2, *p*s > 0.55). Sensitivity and specificity varied across models, ranging from 39-67% and 54-80%, respectively. Sensitivity was highest in the global theta model, while specificity was highest in the theta/gamma ratio model. The best predictors of MCI status were left temporo-parietal theta power and temporal delta power in the EO models (see Figure 4B). The best predictor of HC status was frontal beta2 in the relative power EO model (see Figure 4B).

Relative power EO was the most predictive model for MCI vs AD classification (AUC = 0.67), out-performing 97% of null model iterations (*t*(198) = 29.20, *p* < 0.001). Similar results were found for relative power EC (AUC = 0.64), global theta (AUC = 0.64), absolute power EO (AUC = 0.62) and theta/gamma ratio models (AUC = 0.61), which out-performed >94% of null model iterations (all *t*s > 24, all *p*s < 0.001). Absolute power EC and alpha2/alpha1 ratio models produced weaker results (AUCs ≤ 0.55; *t*s < 9, *p*s < 0.001). Sensitivity and specificity values ranged from 46-73% and 44-74%, respectively. Sensitivity was highest in the alpha2/alpha1 model, while specificity was highest in the relative power EC model. The best predictor of AD status was theta power in left frontal and right parietal regions (see Figure 4C). The best predictor of MCI status was right temporo-parietal beta1 power (see Figure 4C).

#### 3.3.3. MRI models

Data from 38 participants (14 AD, 11 MCI and 13 HC) were excluded due to bad orientation and/or bad quality post-processing.

Figure 5 illustrates ROCs for the MRI models. The AD vs MCI model had perfect predictive accuracy (AUC = 1.00). The AD vs HC model also produced excellent results (AUC = 0.99), while MCI vs HC model performance was moderate (AUC = 0.72). Prediction indices were high for all models, except for MCI vs HC classification, which was only moderately sensitive (i.e. 63%). A large number of predictors (>1000) across multiple brain regions survived the selection threshold in both AD models (see Figure 6A,C). The best predictors of HC status in the MCI vs HC model were the left hippocampus and thalamus (see Figure 6). The best predictors of HC status in the MCI vs HC model were the left hippocampus and thalamus (see Figure 6B). All models out-performed 100% of null model iterations (all *t*s > 46, all *p*s < 0.001).

**Figure 5.**
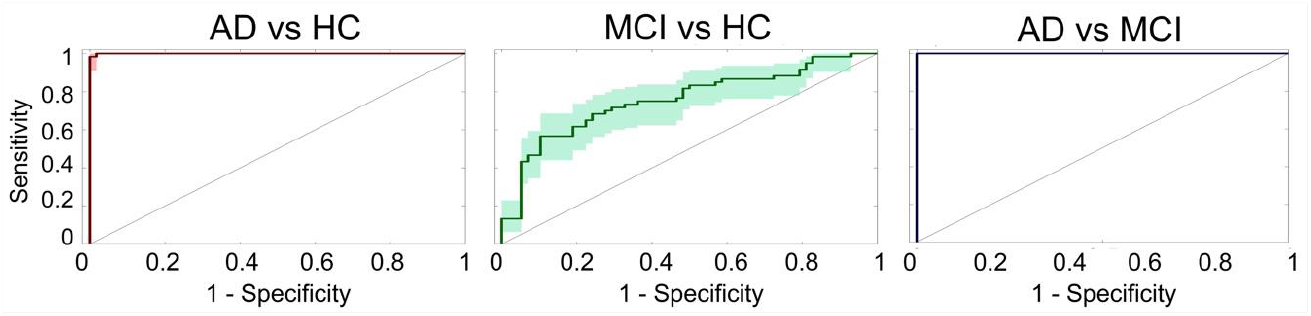
Receiver Operating Curves (ROCs) for MRI actual models with 95% confidence intervals. Curves close to the top left corner (0.0, 1.0) indicate good performance.

**Figure 6.**
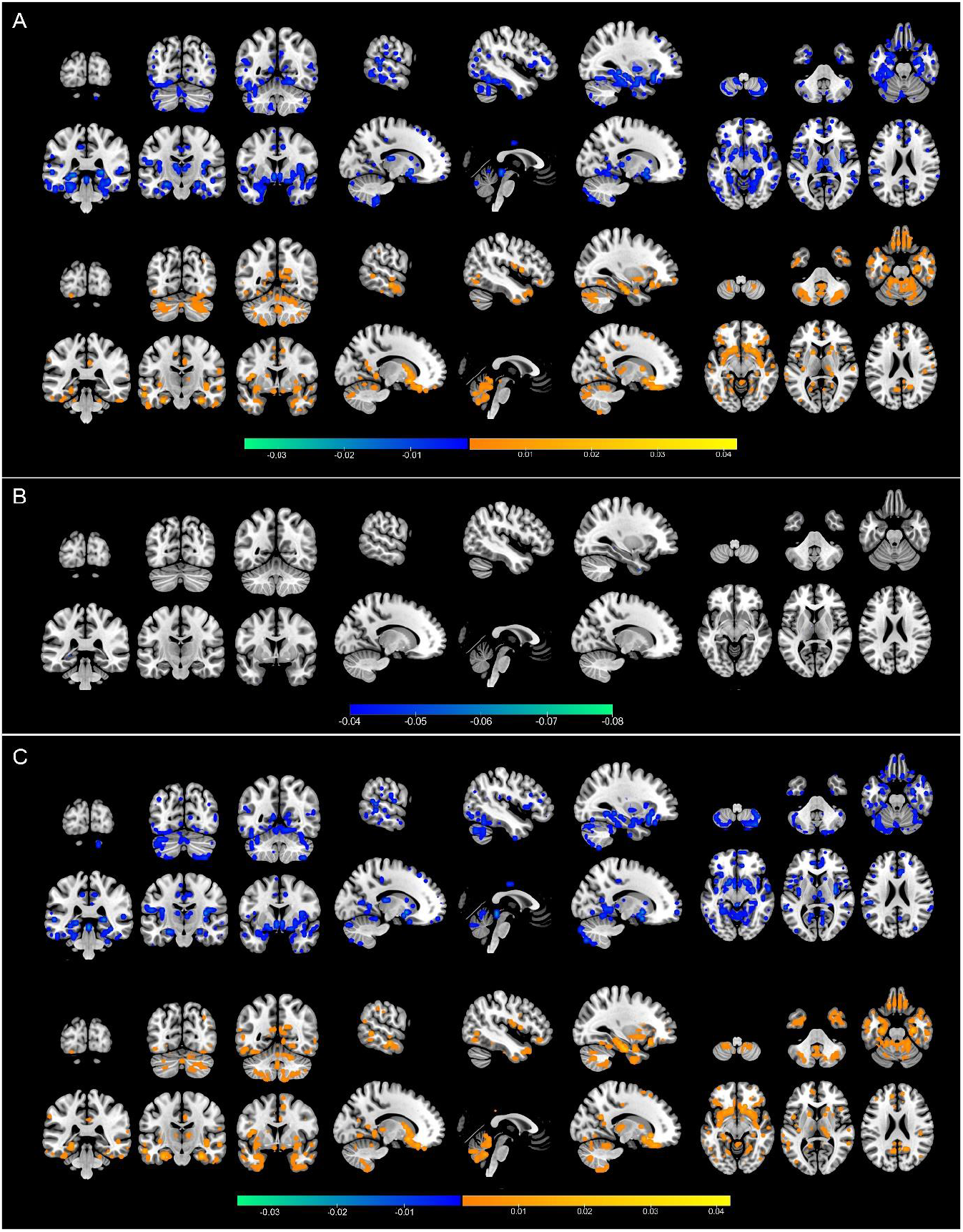
Coronal, sagittal and axial images showing voxels (5 mm^3^) that survived selection frequency for (A) AD vs HC, (B) MCI vs HC and (C) AD vs MCI models. The colour represents beta values ranging from negative (HC/MCI) to positive (MCI/AD).

#### 3.3.4. MMSE+EEG models

Figure 7 illustrates Receiver Operating Curves (ROCs) for the MMSE+EEG models. The AD vs HC model produced the best performance (AUC = 0.98). The next best model was AD vs MCI (AUC = 0.86), followed by the MCI vs HC model (AUC = 0.80). Prediction indices were high for all models (i.e. > 73%). All models significantly out-performed 100% of null model iterations (all *t*s > 91, all *p*s < 0.001). MMSE scores were predictive of HC status in the AD and MCI models, and HC status in the MCI vs HC model. Global theta was predictive of AD status in the HC and MCI models, and MCI status in the MCI vs HC model.

**Figure 7.**
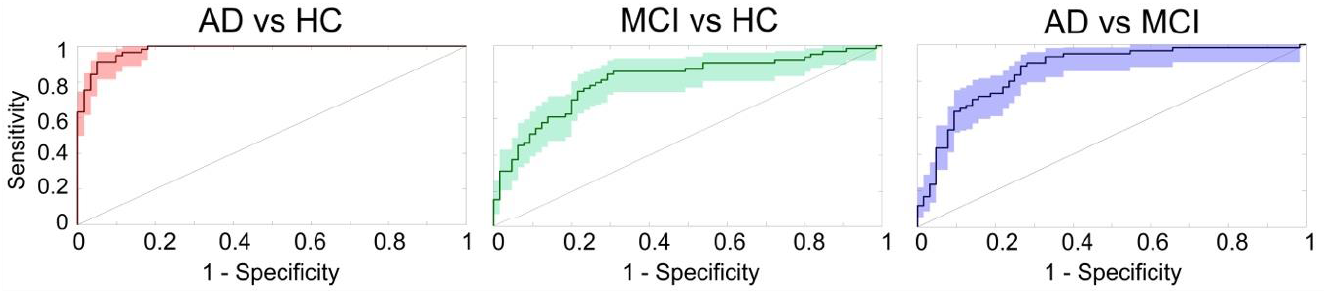
Receiver Operating Curves (ROCs) for MMSE+EEG actual models with 95% confidence intervals. Curves close to the top left corner (0.0, 1.0) indicate good performance.

## 4. Discussion

Early diagnosis of dementia on clinical grounds is challenging. Hence, there is a need for automated tools capable of identifying pathological markers before behavioural symptom onset. The aim of this study was to evaluate the efficacy of resting state EEG in detecting AD and its prodromal stage, MCI. Power spectral analysis was performed on eyes open and eyes closed resting state EEG data from AD, MCI and healthy participants. A machine learning classification analysis was carried out using absolute and relative power across seven frequency bands (delta, theta, low alpha, high alpha, low beta, high beta and gamma), theta/gamma ratio, alpha2/alpha1 ratio and global theta power. We also assessed the utility of a combined behavioural-EEG model using global theta power and MMSE scores. EEG was moderately accurate at classifying AD and HC participants. The best performing models were absolute and relative power and global theta power (AUCs = 0.73-0.76). EEG was less accurate at differentiating AD and MCI participants (highest AUC = 0.67 for relative power EO) and MCI and HC participants (highest AUC = 0.61 for absolute power EO). Overall, EEG models had higher specificity than sensitivity, suggesting a tendency to misidentify patients as being healthy. The combination of global theta power and MMSE scores had no additional value compared to MMSE scores alone (combined AUCs = 0.80-0.98 vs MMSE AUCs = 0.77-0.97).

Our secondary aim was to compare the classification accuracy of resting state EEG with that of structural MRI. Although structural MRI has received considerable attention as a dementia diagnosis tool (e.g. Beheshti et al., 2017), the need for more widely accessible, cost-effective approaches has long been recognised (Jelic et al., 1999). Nonetheless, classification accuracy of EEG and MRI have yet to be investigated in AD and MCI samples. Here, grey matter density estimates were used as inputs for a separate machine learning analysis. MRI was highly accurate at distinguishing AD participants from HC and MCI participants (AUCs = 1.00). On the other hand, classification of MCI and HC participants was only moderately good (AUC = 0.73). Like EEG, specificity was higher than sensitivity, indicating a tendency to under-classify MCI patients. Taken together, our results suggest that structural MRI is a better indicator of AD than resting state EEG. However, both EEG and MRI have limited sensitivity in distinguishing MCI from healthy aging.

Our EEG results are largely consistent with previous literature. Babilioni and colleagues reported similar accuracy rates of up to 82% for classification of AD and HC participants using absolute power EC data (Babiloni et al., 2016; Triggiani et al., 2017), while Rodriguez *et al*. (1998) reported 77% accuracy for relative power EC. MCI classification accuracy was also similar to that reported by Musaeus *et al*. (2018) for absolute power EC and EO models, which ranged between 60-63%. Accuracy for the MCI vs AD classification model was lower than previously reported by Hatz and colleagues (AUC = 0.77) (2015). This is likely due to differences in the type of EEG data used (i.e. theta power connectivity by Hatz *et al*.) but may also have been influenced by small sample size (which can inflate classification accuracy; Mateos-Pérez et al., 2018). In addition, we found global theta power to be a better classifier of AD and HC participants than Musaeus and colleagues (2018), who reported 54% accuracy with a model combining global theta power and left alpha coherence.

Theta power in frontal, temporal and parietal regions emerged as an important EEG characteristic of AD and MCI in this study. Theta power was highest in the AD group, followed by the MCI group and the HC group. The opposite pattern was found for beta power, such that beta was highest in HC participants and lowest in AD participants. MCI and HC participants were discriminated by (high) beta power in frontal regions, while AD participants were distinguished from MCI and HC participants by (low) beta in temporo-parietal regions. These findings are in keeping with a global shift towards lower frequencies typically observed AD and MCI (Babiloni et al., 2006; Musaeus et al., 2018). Increased alpha2/alpha1 ratio power in left temporo-parietal regions was identified as marker of healthy aging. Importantly, alpha2/alpha1 ratio power was a predictor of HC status when compared with AD only (i.e., not with MCI), suggesting that alpha may be more useful as a late-stage disease marker. Conversely, temporal delta power distinguished MCI participants from healthy older adults, but not from AD participants, indicating that delta may be a useful indicator of early impairment. This is consistent with research by Morretti and colleagues, who proposed that delta and theta power are higher in MCI patients who will convert to AD, relative to those who will not (Moretti et al., 2009; Moretti et al., 2012).

MRI results were also in accordance with previous research (Beheshti et al., 2017). AD classification models were highly accurate, as expected. AD was associated with widespread grey matter atrophy, including frontal, temporal, parietal and occipital lobes, as well as the cerebellum. MRI predictors were more left than right lateralised overall. This is consistent with work showing that grey matter loss in AD is faster in the left hemisphere than in the right (Thompson et al., 2003). Classification of MCI and HC participants was less accurate, but still moderately good. The left hippocampus and thalamus were identified as important predictors in the MCI vs HC model. Temporal lobe atrophy (including the hippocampus) is a well-established biomarker of neurodegeneration in dementia (Jack Jr et al., 2018). Atrophy of the thalamus has also been strongly linked to AD and MCI (de Jong et al., 2008; Hahn, Lee, Won, Joo, & Lim, 2016). Specifically, Hahn and colleagues showed that left thalamic volumes were significantly smaller in MCI patients compared to healthy control participants.

A key finding from this study is that neither resting state EEG nor structural MRI were sufficiently sensitive to distinguish MCI from healthy aging. None of the measures used (including MMSE) achieved sensitivity above 70%. This is an important finding given that early-identification is a key priority in dementia research, particularly for low- and middle-income countries where the prevalence of dementia is expected to increase dramatically over the next 30 years (Prince et al., 2015). There are a number of factors that can hinder MCI classification. Firstly, the pathological changes that occur in dementia can emerge up to 20 years before any symptoms appear; thus, it is possible that some participants included as HCs were in a pre-clinical stage of dementia (Guo et al., 2014). Secondly, the heterogeneity of MCI can lead to considerable diagnostic variability among clinicians (Duara et al., 2010). This imposes a limit on the accuracy that can realistically be achieved using classification methods. Thirdly, MCI is an unstable condition (Petersen et al., 2014). Thus, it is unclear whether (and when) individuals with MCI may revert to healthy status.

This study had multiple positive aspects, including extensive clinical and neuropsychological profiling of participants, matched groups (for age, sex and education) and a good sample size relative to earlier studies (e.g. Ahmadlou, Adeli, & Adeli, 2010; Bertè, Lamponi, Calabrò, & Bramanti, 2014; Kashefpoor, Rabbani, Barekatain, 2016; McBride et al., 2015; Musaeus et al., 2018). Additionally, our MCI group included only participants with amnestic MCI sub-type, which is more closely linked with AD than non-amnestic MCI (Csukly et al., 2016). This study was also the first to investigate resting state EEG and structural MRI markers of AD and MCI in participants who underwent the sample neuropsychological evaluation and testing. Finally, we employed a machine learning approach with penalized regression and cross-validation, thereby maximising the reliability of our findings (Mateos-Pérez et al., 2018). The main limitation of this study was the lack of longitudinal data. Without these data, we cannot rule out the possibility that some participants’ diagnostic status changed after the test date. A second limitation was that our machine learning analysis did not include an external validation set, thereby limiting the generalisability of our findings (Mateos-Pérez et al., 2018).

Future work should focus on longitudinal data collection in order to more accurately assess the sensitivity and specificity of resting state EEG markers for predicting AD and MCI (Moretti et al., 2012). Future research should also assess the utility of task-based EEG data for dementia diagnosis. One recent study by Porcaro, Balsters, Mantini, Robertson and Wenderoth (2019) suggested the P3b event-related potential (ERP) as an early marker of memory dysfunction in older adults. However, more work is needed to support the use of ERP components in AD and MCI classification. An additional recommendation for future work is to evaluate resting state EEG as a tool to differentiate AD from other neurological disorders (e.g., Parkinson’s disease). Identifying EEG markers specific to AD or MCI may be helpful at early stages of the disease when symptoms overlap (Cassani et al., 2018). Some research has been carried out in this area to date with vascular dementia and frontotemporal dementia, however, results are limited (see Neto, Biessmann, Aurlien, Nordby, & Eichele, 2016; Nishida et al., 2011).

The need for low-cost, accessible tools to aid dementia diagnosis is particularly pertinent today, given the projected increase in dementia incidence in low- and middle-income countries. EEG holds great promise as one such tool. In the current study, we evaluated the utility of resting state EEG for classification of AD and MCI, relative to healthy controls, in an older adult sample. We also assessed the performance of combined behavioural-EEG and structural MRI classification models. Structural MRI out-performed resting state EEG for AD classification. However, both EEG and MRI had limited utility in detecting MCI. All models (including MMSE scores) had a tendency to under-classify MCI, and none were sufficiently sensitive (i.e. <70%). Findings indicate that resting state EEG does not appear to be a suitable tool for classifying AD. However, EEG classification accuracy was comparable to structural MRI when distinguishing MCI from healthy aging, although neither were sufficiently accurate to have clinical utility. Suggestions for future work include prospective studies and MCI classification using task-based EEG.

## Supporting information

Supplementary model features

## Acknowledgements

The data collection was partially supported by the Turkish National Science and Research Council (TUBITAK, Grant number: 112S459) and the Dokuz Eylul University Scientific Research Projects (Grant number: 2018.KB.SAG.084). The funding agencies had no involvement in the conduct of the research or preparation of the article. Francesca Farina and Rory Boyle are supported by the Irish Research Council (IRC) under project numbers EPSPD/2017/110 and EPSPG/2017/277, respectively. Hanni Kiiski was also supported by the IRC (grant number: GOIPD/2015/777). Robert Whelan is supported by Science Foundation Ireland (project number: 16/ERCD/3797).

## Supplementary Materials

**Figure S1.**
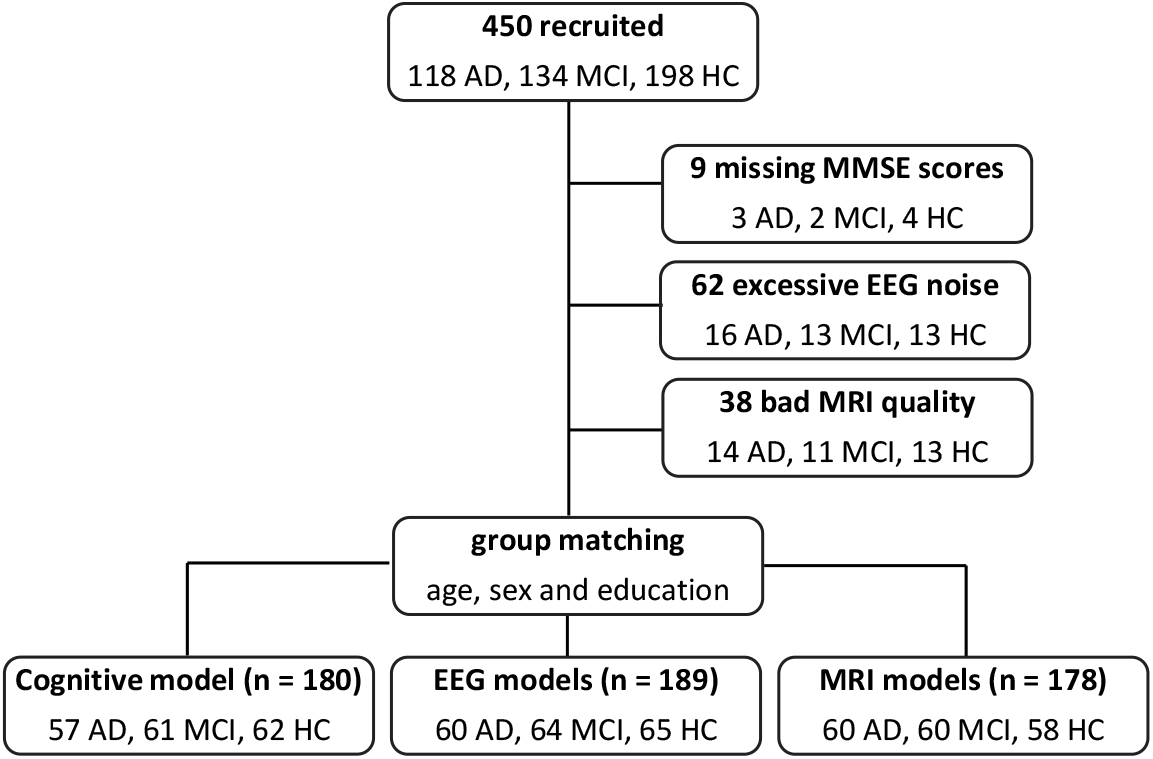
Flow chart of participant inclusions and exclusions.

**Figure S2.**
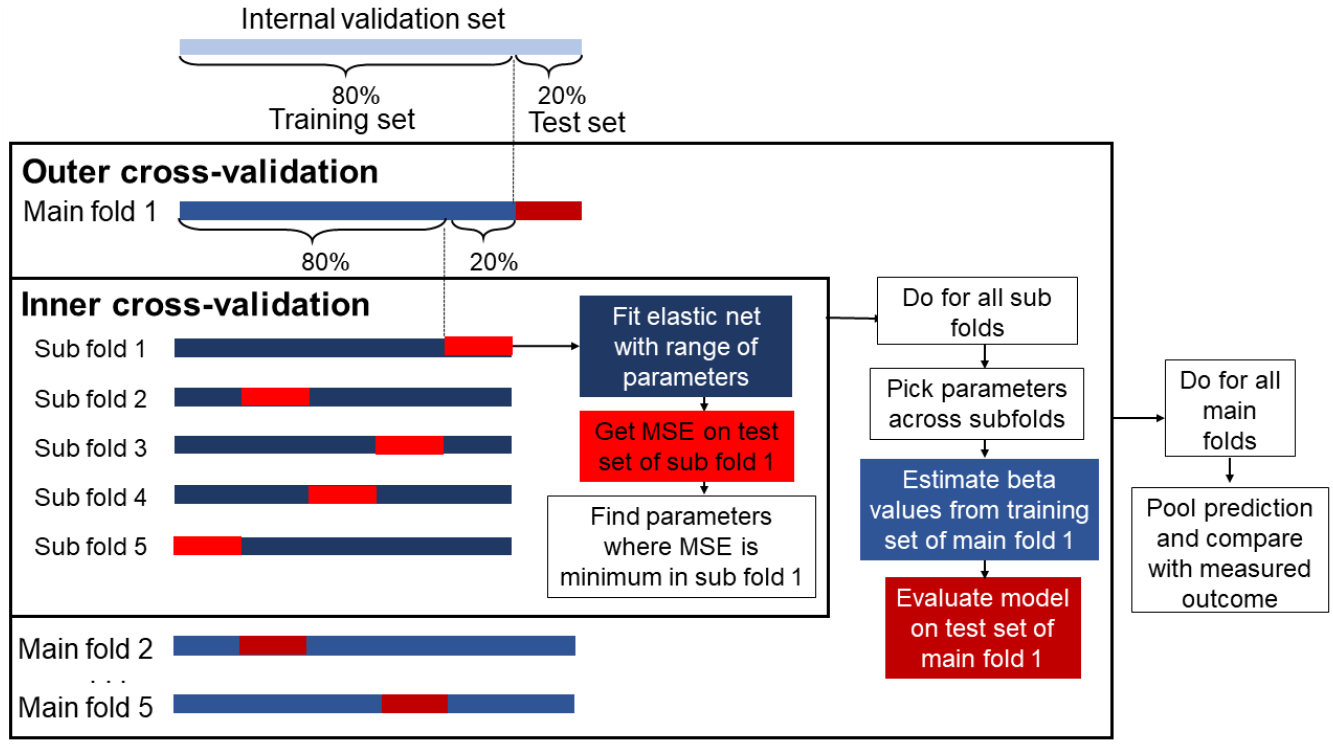
Procedure for the nested cross-validation. MSE=mean squared error (adapted from Rueda-Delgado et al., 2019).

## Notes

**Conflict of interest:** none declared

## References

Ahmadlou, M., Adeli, H., & Adeli, A. (2010). New diagnostic EEG markers of the Alzheimer’s disease using visibility graph. Journal of neural transmission, 117(9), 1099–1109.

Albert, M. S., DeKosky, S. T., Dickson, D., Dubois, B., Feldman, H. H., Fox, N. C., … & Snyder, P. J. (2011). The diagnosis of mild cognitive impairment due to Alzheimer’s disease: Recommendations from the National Institute on Aging-Alzheimer’s Association workgroups on diagnostic guidelines for Alzheimer’s disease. Alzheimer’s & dementia, 7(3), 270–279.

Apostolova L. G. (2016). Alzheimer Disease. Continuum (Minneapolis, Minn.), 22(2 Dementia), 419–434.

Babiloni, C., Binetti, G., Cassetta, E., Dal Forno, G., Del Percio, C., Ferreri, F., … Lanuzza, B. J. C. n. (2006). Sources of cortical rhythms change as a function of cognitive impairment in pathological aging: a multicenter study. Clinical neurophysiology, 117(2), 252–268.

Babiloni, C., Triggiani, A. I., Lizio, R., Cordone, S., Tattoli, G., Bevilacqua, V., … Gesualdo, L. (2016). Classification of single normal and Alzheimer’s disease individuals from cortical sources of resting state EEG rhythms. Frontiers in neuroscience, 10, 47.

Barry, R. J., & De Blasio, F. M. (2017). EEG differences between eyes-closed and eyes-open resting remain in healthy ageing. Biological Psychology, 129, 293–304.

Beheshti, I., Demirel, H., Matsuda, H., & Alzheimer’s disease Neuroimaging Initiative. (2017). Classification of Alzheimer’s disease and prediction of mild cognitive impairment-to-Alzheimer’s conversion from structural magnetic resource imaging using feature ranking and a genetic algorithm. Computers in biology medicine, 83, 109–119.

Bertè, F., Lamponi, G., Calabrò, R. S., & Bramanti, P. (2014). Elman neural network for the early identification of cognitive impairment in Alzheimer’s disease. Functional neurology, 29(1), 57.

Cassani, R., Estarellas, M., San-Martin, R., Fraga, F. J., & Falk, T. H. (2018). Systematic review on resting-state EEG for Alzheimer’s disease diagnosis and progression assessment. Disease markers, 2018.

Csukly, G., Sirály, E., Fodor, Z., Horváth, A., Salacz, P., Hidasi, Z., … Szabó, Á. (2016). The Differentiation of Amnestic Type MCI from the Non-Amnestic Types by Structural MRI. Frontiers in aging neuroscience, 8, 52–52.

Dauwels, J., Vialatte, F., & Cichocki, A. (2010). Diagnosis of Alzheimer’s disease from EEG signals: where are we standing? Current Alzheimer Research, 7(6), 487–505.

de Jong, L. W., van der Hiele, K., Veer, I. M., Houwing, J. J., Westendorp, R. G. J., Bollen, E. L. E. M., … van der Grond, J. (2008). Strongly reduced volumes of putamen and thalamus in Alzheimer’s disease: an MRI study. Brain, 131 (Pt 12), 3277–3285.

Delorme, A., & Makeig, S. (2004). EEGLAB: an open source toolbox for analysis of single-trial EEG dynamics including independent component analysis. Journal of neuroscience methods, 134(1), 9–21.

Duara, R., Loewenstein, D. A., Greig, M., Acevedo, A., Potter, E., Appel, J., … Potter, H. (2010). Reliability and validity of an algorithm for the diagnosis of normal cognition, mild cognitive impairment, and dementia: implications for multicenter research studies. The American journal of geriatric psychiatry: official journal of the American Association for Geriatric Psychiatry, 18(4), 363–370.

Dubois, B., Feldman, H. H., Jacova, C., DeKosky, S. T., Barberger-Gateau, P., Cummings, J., … Jicha, G. J. (2007). Research criteria for the diagnosis of Alzheimer’s disease: revising the NINCDS-ADRDA criteria. The Lancet Neurology, 6(8), 734–746.

Folstein, M. F., Folstein, S. E., & McHugh, P. R. (1975). “Mini-mental state”: a practical method for grading the cognitive state of patients for the clinician. Journal of Psychiatric Research, 12(3), 189–198.

Guo, Y., Zhang, Z., Zhou, B., Wang, P., Yao, H., Yuan, M., … Zhang, X. (2014). Grey-matter volume as a potential feature for the classification of Alzheimer’s disease and mild cognitive impairment: an exploratory study. Neuroscience bulletin, 30(3), 477–489.

Hahn, C., Lee, C.-U., Won, W. Y., Joo, S.-H., & Lim, H. K. (2016). Thalamic shape and cognitive performance in amnestic mild cognitive impairment. Psychiatry investigation, 13(5), 504.

Hatz, F., Hardmeier, M., Benz, N., Ehrensperger, M., Gschwandtner, U., Rüegg, S., … & Fuhr, P. (2015). Microstate connectivity alterations in patients with early Alzheimer’s disease. Alzheimer’s research & therapy, 7(1), 78.

Hughes, C. P., Berg, L., Danziger, W., Coben, L. A., & Martin, R. L. (1982). A new clinical scale for the staging of dementia. The British journal of psychiatry, 140(6), 566–572.

Jack, C. R., Bennett, D. A., Blennow, K., Carrillo, M. C., Dunn, B., Haeberlein, S. B., … & Liu, E. (2018). NIA-AA Research Framework: Toward a biological definition of Alzheimer’s disease. Alzheimer’s & Dementia, 14(4), 535–562.

Jelic, V., Wahlund, L., Almkvist, O., Johansson, S., Shigeta, M., Winblad, B., & Nordberg, A. (1999). Diagnostic accuracies of quantitative EEG and PET in mild Alzheimer’s disease. Alzheimer’s Reports, 2(5), 291–298.

Jicha, G. A., Parisi, J. E., Dickson, D. W., Johnson, K., Cha, R., Ivnik, R. J., … Braak, H. J. (2006). Neuropathologic outcome of mild cognitive impairment following progression to clinical dementia. Archives of neurology, 63(5), 674–681.

Kaplan, E., Goodglass, H., & Weintraub, S. J. A., TX: Pro-Ed. (2001). Boston naming test-2 (BNT-2).

Kashefpoor, M., Rabbani, H., & Barekatain, M. (2016). Automatic diagnosis of mild cognitive impairment using electroencephalogram spectral features. Journal of medical signals and sensors, 6(1), 25.

Kiiski, H., Jollans, L., Donnchadha, S. Ó., Nolan, H., Lonergan, R., Kelly, S., … Burke, T. J. B. t. (2018). Machine learning EEG to predict cognitive functioning and processing speed over a 2-year period in multiple sclerosis patients and controls. Brain Topography, 31(3), 346–363.

Lehmann, C., Koenig, T., Jelic, V., Prichep, L., John, R. E., Wahlund, L. O., … & Dierks, T. (2007). Application and comparison of classification algorithms for recognition of Alzheimer’s disease in electrical brain activity (EEG). Journal of neuroscience methods, 161(2), 342–350.

Lizio, R., Del Percio, C., Marzano, N., Soricelli, A., Yener, G. G., Başar, E., … Ferri, R. (2016). Neurophysiological assessment of Alzheimer’s disease individuals by a single electroencephalographic marker. Journal of Alzheimer’s disease, 49(1), 159–177.

Mack, W. J., Freed, D. M., Williams, B. W., & Henderson, V. W. (1992). Boston Naming Test: shortened versions for use in Alzheimer’s disease. Journal of gerontology, 47(3), 154–158.

Mateos-Pérez, J. M., Dadar, M., Lacalle-Aurioles, M., Iturria-Medina, Y., Zeighami, Y., & Evans, A. C. (2018). Structural neuroimaging as clinical predictor: A review of machine learning applications. NeuroImage: Clinical, 20, 506–522.

McBride, J. C., Zhao, X., Munro, N. B., Jicha, G. A., Schmitt, F. A., Kryscio, R. J., … & Jiang, Y. (2015). Sugihara causality analysis of scalp EEG for detection of early Alzheimer’s disease. NeuroImage: Clinical, 7, 258–265.

McKhann, G. M., Knopman, D. S., Chertkow, H., Hyman, B. T., Jack Jr, C. R., Kawas, C. H., … & Mohs, R. C. (2011). The diagnosis of dementia due to Alzheimer’s disease: Recommendations from the National Institute on Aging-Alzheimer’s Association workgroups on diagnostic guidelines for Alzheimer’s disease. Alzheimer’s & dementia, 7(3), 263–269.

Michaud, T. L., Su, D., Siahpush, M., & Murman, D. L. (2017). The risk of incident mild cognitive impairment and progression to dementia considering mild cognitive impairment subtypes. Dementia and geriatric cognitive disorders extra, 7(1), 15–29.

Moretti, D. V., Pievani, M., Fracassi, C., Binetti, G., Rosini, S., Geroldi, C., … & Frisoni, G. B. (2009). Increase of theta/gamma and alpha3/alpha2 ratio is associated with amygdalo-hippocampal complex atrophy. Journal of Alzheimer’s disease, 17(2), 349–357.

Moretti, D. V., Prestia, A., Fracassi, C., Binetti, G., Zanetti, O., & Frisoni, G. B. (2012). Specific EEG changes associated with atrophy of hippocampus in subjects with mild cognitive impairment and Alzheimer’s disease. International Journal of Alzheimer’s disease, 2012.

Musaeus, C. S., Engedal, K., Hgh, P., Jelic, V., Mrup, M., Naik, M., … & Bo Andersen, B. (2018). EEG theta power is an early marker of cognitive decline in dementia due to Alzheimer’s disease. Journal of Alzheimer’s disease, (Preprint), 1–13.

Nasreddine, Z. S., Phillips, N. A., Bédirian, V., Charbonneau, S., Whitehead, V., Collin, I., … & Chertkow, H. (2005). The Montreal Cognitive Assessment, MoCA: a brief screening tool for mild cognitive impairment. Journal of the American Geriatrics Society, 53(4), 695–699.

Neto, E., Biessmann, F., Aurlien, H., Nordby, H., & Eichele, T. (2016). Regularized linear discriminant analysis of EEG features in dementia patients. Frontiers in aging neuroscience, 8, 273.

Nishida, K., Yoshimura, M., Isotani, T., Yoshida, T., Kitaura, Y., Saito, A., … Suwa, A. (2011). Differences in quantitative EEG between frontotemporal dementia and Alzheimer’s disease as revealed by LORETA. Clinical Neurophysiology, 122(9), 1718–1725.

Nolan, H., Whelan, R., & Reilly, R. B. (2010). FASTER: fully automated statistical thresholding for EEG artifact rejection. Journal of neuroscience methods, 192(1), 152–162.

Ogbole, G. I., Adeyomoye, A. O., Badu-Peprah, A., Mensah, Y., & Nzeh, D. A. (2018). Survey of magnetic resonance imaging availability in West Africa. Pan African Medical Journal, 30(1).

Öktem, Ö. (1992). Sözel Bellek Süreçleri Testi, Bir ön çalışma. Nöropsikiyatri Arşivi, 29.

Olsson, B., Lautner, R., Andreasson, U., Öhrfelt, A., Portelius, E., Bjerke, M., … & Wu, E. (2016). CSF and blood biomarkers for the diagnosis of Alzheimer’s disease: a systematic review and meta-analysis. The Lancet Neurology, 15(7), 673–684.

Park, M., & Moon, W. J. (2016). Structural MR imaging in the diagnosis of Alzheimer’s disease and other neurodegenerative dementia: current imaging approach and future perspectives. Korean journal of radiology, 17(6), 827–845.

Parra, M. A. (2014). Overcoming barriers in cognitive assessment of Alzheimer’s disease. Dementia & neuropsychologia, 8(2), 95–98.

Petersen, R. C., Caracciolo, B., Brayne, C., Gauthier, S., Jelic, V., & Fratiglioni, L. (2014). Mild cognitive impairment: a concept in evolution. Journal of internal medicine, 275(3), 214–228.

Petersen, R. C., & Negash, S. (2008). Mild cognitive impairment: an overview. CNS spectrums, 13(1), 45–53.

Porcaro, C., Balsters, J. H., Mantini, D., Robertson, I. H., & Wenderoth, N. (2019). P3b amplitude as a signature of cognitive decline in the older population: An EEG study enhanced by Functional Source Separation. NeuroImage, 184, 535–546.

Prince, M., Wimo, A., Guerchet, M., Ali, G., Wu, Y., & Prina, M. (2015). World Alzheimer Report 2015. The Global Impact of Dementia. Alzheimer’s disease International, London.

Raamana, P. R., Weiner, M. W., Wang, L., Beg, M. F., & Alzheimer’s disease Neuroimaging Initiative. (2015). Thickness network features for prognostic applications in dementia. Neurobiology of aging, 36, S91–S102.

Rodriguez, G., Nobili, F., Rocca, G., De Carli, F., Gianelli, M. V., & Rosadini, G. (1998). Quantitative electroencephalography and regional cerebral blood flow: discriminant analysis between Alzheimer’s patients and healthy controls. Dementia and geriatric cognitive disorders, 9(5), 274–283.

Rossini, P. M., Buscema, M., Capriotti, M., Grossi, E., Rodriguez, G., Del Percio, C., & Babiloni, C. (2008). Is it possible to automatically distinguish resting EEG data of normal elderly vs. mild cognitive impairment subjects with high degree of accuracy? Clinical Neurophysiology, 119(7), 1534–1545.

Shah, H., Albanese, E., Duggan, C., Rudan, I., Langa, K. M., Carrillo, M. C., … Rossor, M. J. T. L. N. (2016). Research priorities to reduce the global burden of dementia by 2025. The Lancet Neurology, 15(12), 1285–1294.

Thompson, P. M., Hayashi, K. M., De Zubicaray, G., Janke, A. L., Rose, S. E., Semple, J., … Doddrell, D. M. (2003). Dynamics of gray matter loss in Alzheimer’s disease. Journal of neuroscience, 23(3), 994–1005.

Triggiani, A. I., Bevilacqua, V., Brunetti, A., Lizio, R., Tattoli, G., Cassano, F., … Gesualdo, L. (2017). Classification of healthy subjects and Alzheimer’s disease patients with dementia from cortical sources of resting state EEG rhythms: a study using artificial neural networks. Frontiers in neuroscience, 10, 604.

Tsolaki, A., Kazis, D., Kompatsiaris, I., Kosmidou, V., & Tsolaki, M. (2014). Electroencephalogram and Alzheimer’s disease: clinical and research approaches. International journal of Alzheimer’s disease, 2014, 349249–349249.

Tumac, A. (1997). Effects of age and education to performance in some frontal lobe tests in normal subjects. Master Thesis, Istanbul University Institute of Social Sciences Department of Psychology, Istanbul, 1997 (Turkish).

Varghese, T., Sheelakumari, R., James, J. S., & Mathuranath, P. S. (2013). A review of neuroimaging biomarkers of Alzheimer’s disease. Neurology Asia, 18(3), 239.

Varoquaux, G., Raamana, P. R., Engemann, D. A., Hoyos-Idrobo, A., Schwartz, Y., & Thirion, B. (2017). Assessing and tuning brain decoders: cross-validation, caveats, and guidelines. NeuroImage, 145, 166–179.

Wechsler, D. (1987). Wechsler memory scale-revised. Psychological Corporation.

Weller, J., & Budson, A. (2018). Current understanding of Alzheimer’s disease diagnosis and treatment. F1000Research, 7.

Yesavage, J. A., Brink, T. L., Rose, T. L., Lum, O., Huang, V., Adey, M., & Leirer, V. O. (1982). Development and validation of a geriatric depression screening scale: a preliminary report. Journal of psychiatric research, 17(1), 37–49.

